# Heparin-modified alginate microspheres enhance neovessel formation in hiPSC-derived endothelial cells and heterocellular *in vitro* models by controlled release of VEGF

**DOI:** 10.1101/2021.01.08.425908

**Authors:** Fabiola Munarin, Carly Kabelac, Kareen L.K. Coulombe

## Abstract

A formidable challenge in regenerative medicine is the development of stable microvascular networks to restore adequate blood flow or to sustain graft viability and long-term function in implanted or ischemic tissues. In this work, we develop a biomimetic approach to increase the binding affinity of the extracellular matrix for the class of heparin-binding growth factors to localize and control the release of proangiogenic cues while maintaining their bioactivity. Sulfate and heparin moieties are covalently coupled to alginate, and alginate microspheres are produced and used as local delivery depots for vascular endothelial growth factor (VEGF). Release of VEGF from sulfate-alginate and heparin-alginate bulk hydrogels and microspheres was sustained over 14 days. *In vitro* evaluation with human induced pluripotent stem cell (hiPSC)-derived endothelial cells and aortic ring assay in a chemically-defined hydrogel demonstrates development of primitive three-dimensional vessel-like networks in the presence of VEGF released from the chemically modified alginate microspheres. Furthermore, our results suggest that the sulfate groups available on the chemically modified alginate microspheres promote some new vessel formation even in VEGF-free samples. Based on this evidence, we conclude that sulfate- and heparin-alginate hydrogels are adaptive and bioactive delivery systems for revascularization therapy and translational vascular tissue engineering.

## 1. Introduction

Revascularization of damaged tissues and vascular integration of engineered tissues with the host blood supply remain crucial challenges for tissue regeneration and regenerative medicine applications. Unless a functional vascular network can rapidly perfuse the injured or implanted tissues, cells will necrose due to the insufficient exchange of oxygen and nutrients [1, 2]. Angiogenesis is a crucial process in wound healing and tissue revascularization that begins with sprouting of new capillaries from the existing vasculature to develop into a functional vascular bed that nourishes and supports the tissue repair process. This process is regulated by gradients of endogenous cytokines including VEGF, bFGF, angiopoietin, and others [3]. The main factors believed to limit the efficacy of exogenous growth factors delivered *in vivo* are primarily their short half-life, followed by rapid convection/diffusion away from the target site and reduced bioactivity [4]. To compensate for the rapid need of microvasculature, biomaterials-based regeneration strategies have been developed that encompass acellular angiogenic treatments and tissue engineering approaches [5-11].

In this work we propose an acellular approach to stimulate neovascularization in ischemic tissues that consists of amplifying the native host response by supplying angiogenic growth factors in the local microenvironment to promote new vessel development [12, 13]. We and others have previously used alginate hydrogels and microspheres for the release of growth factors, due to the minimal processing, rapid encapsulation of bioactive molecules, and high compatibility with cells and tissues [12-19]. However, immobilized growth factors are only retained in alginate hydrogels by relatively weak ionic interactions, and thus, in the presence of diffusion, relatively short release kinetics of growth factors are obtained from alginate microspheres [20]. Controlling the release of growth factors from alginate hydrogels up to 2 weeks would enable us to design release profiles that better match the kinetics of angiogenesis and wound healing and enable more precise orchestration of the therapeutic release, particularly when using cocktails of multiple proteins. In this study, we evaluate the release of a crucial signaling molecule, vascular endothelial growth factor (VEGF), from sulfate- and heparin-conjugated alginate hydrogels, and we perform assays in clinically relevant cell models. Biomaterials screening for vascular biology is often performed with human umbilical vein endothelial cells (HUVECs) that are very robust, widely used experimentally, however they lack translational potential for cell-based approaches. The use of a well-defined source of human induced pluripotent stem cell-derived endothelial cells (hiPSC-ECs) allows us to assess angiogenesis and blood vessel formation with a more clinically relevant approach for cell-based tissue engineering applications.

## 2. Materials and Methods

### 2.1 Materials

The materials required for alginate chemical modifications and microspheres production included alginate (180947, Sigma Aldrich), CaCl_2_ (Fisher, S25223), NaCl (Fisher, BP358-10), NaOH (Sigma Aldrich 221465), ethylenediamine (EDA, Sigma Aldrich E1521), 1-ethyl-3-(3-dimethylaminopropyl)-carbodiimide hydrochloride (EDC-HCl, Sigma Aldrich, 8510070025), chlorosulfonic acid (Sigma 571024), formamide (Sigma Aldrich F9037), heparin (Sigma, H5515), acetone (Fisher RSOA0010500) and 2-morpholinoethane sulfonic acid (Sigma M3671). For in vitro and ex vivo assays, VEGF (Life Technologies, PHC9391) was reconstituted in 0.1% bovine serum albumin (BSA) and used without further modifications. iCell® Endothelial Cells (Cellular Dynamics, Inc.) were cultured with Lonza EGM-2 BulletKit (CC3156).

### 2.2 Synthesis of sulfated alginate (S-Alg)

Alginate sulfation was performed following published protocols, with slight modifications [21]. Alginate was first hydrolyzed by lowering the pH of a 2% (w/v) aqueous solution to 5.6 with NaOH, followed by heating the solution at 95°C for 1 hour. The hydrolyzed alginate was dialyzed against 1M NaCl followed by deionized water for 24 hours before being lyophilized. Chlorosulfonic acid was added dropwise into 4 ml of formamide to a final concentration of 1.5% (w/v), to yield approximately 0.05 sulfate per alginate monosaccharide [22]. The hydrolyzed alginate powder (10 0mg) was added to the mixture, and agitated at 60°C. After 4 hours, the sulfated alginate was precipitated by the addition of acetone (18 mL) and centrifugation at 5000 rpm for 6 minutes. The precipitate was re-dissolved in water and the pH was adjusted to 7.0 using NaOH 5M. The pH of S-Alg was further lowered to 3 to de-ionize the solution. To remove excess ions and formamide, S-Alg was dialyzed against 100 mM NaCl and deionized water for 24 hours. After dialysis, the pH was raised back to 7 and the S-Alg solution was lyophilized and stored at − 20°C until later use. Fourier transform infrared (FT-IR) spectroscopy was used to detect changes in the chemically modified alginate spectra in the IR region of 4000 cm^-1^ − 400 cm^-1^, with 4 cm^-1^ resolution using a Bruker ALPHA spectrometer.

### 2.3 Synthesis of heparin-conjugated alginate (H-Alg)

Heparin conjugation with alginate was obtained by covalent binding following published protocols [23] with slight modifications. Briefly, alginate 1% (w/v) was dissolved in 0.05M 2-morpholinoethane sulfonic acid (MES buffer, pH 5.4) at RT overnight. In order to activate approximately 10% of the carboxylic groups in alginate [24], 100 mg of EDC and 60 mg of NHS were added per gram of alginate. After 15 minutes, EDA (3.1×10^−2^ M) was added and allowed to react for 4 hours at RT. Simultaneously, heparin (2% (w/v) in 0.05M MES buffer) was mixed with EDC and NHS according to a mole ratio of 0.40:0.24:1.00 (mol_EDC_: mol_NHS_: mol_hep COOH_) and it was gently stirred for 15 minutes at room temperature before being added to the alginate EDA mixture. This covalent binding reaction was carried on for 24 hours at room temperature. In order to remove unreacted EDC and NHS, the resulting polymer was dialyzed for 24 hours against 1M NaCl followed by deionized water for 3 days with frequent changes of the dialysis solutions. The dialyzed H-Alg solution was then lyophilized and kept frozen (−20°C) until later use. Toluidine blue assay was performed to measure the amount of heparin bound to alginate after chemical modification, as follows. A 0.04% (w/v) solution of toluidine blue was prepared in 0.01 M HCl/ 0.2% NaCl. 50 µL of test sample (H-Alg) was added to 50 µL of toluidine blue solution and was incubated at 37° C for 4 h. The formed heparin-toluidine blue precipitate was centrifuged at 3500 rpm for 10 minutes, then rinsed with 0.01 M HCl/ 0.2% NaCl and dissolved in 200 µL of 80% ethanol/ 0.1 M NaOH. Absorbance was measured at 530 nm by use of a Cytation™ 3 Plate Reader (Biotek). For standard curve preparation, aqueous solution with known heparin concentrations in the range of 0 to 2.5 mg/mL were used.

### 2.4 Production of alginate bulk hydrogels and microspheres

1% (w/v) unmodified alginate (Alg) was mixed with 10 mg H-Alg/g of Alg or 5 mg S-Alg/g of Alg and (1) placed in the wells of a 24 wells culture plate (0.5 mL/well), frozen overnight at −20°C, after which 100 µL of 0.15M CaCl_2_ solution was added at 4°C for 3h to allow for uniform internal gelation or (2) sprayed through a VarJ30 Bead Generator (Nisco, Switzerland) into a 0.15M calcium chloride gelling bath (10 mL) for microspheres production. Nitrogen pressure of the Bead Generator was set to 90 mbar, nozzle size to 0.35 mm and velocity of extrusion to 18 ml/h. After preparation, microspheres were filtered through a 40 µm porosity strainer, and rinsed with 5 mL of PBS. Protein-loaded hydrogels and microspheres were obtained by mixing vascular endothelial growth factor (VEGF, 0.5-4 µg/mL) with Alg, H-Alg and S-Alg prior to gelling.

Images of the microspheres were captured after preparation by Nikon Eclipse Ti-E inverted microscope, and the diameters of 3 different batches of microspheres were measured for each formulations with the open source software Fiji [25].

### 2.5 Release kinetics of vascular endothelial growth factor (VEGF) from alginate bulk hydrogels and microspheres

VEGF was immobilized into Alg, S-Alg and H-Alg hydrogels and microspheres and used as a model growth factor to study the release profiles of the different formulations. Unmodified and chemically-modified alginate hydrogels were produced by mixing VEGF (1 µg/mL and 4 µg/mL in bulk hydrogels and microspheres, respectively) with alginate, heparin-alginate or sulfate-alginate solutions. To determine VEGF release kinetics, the bulk hydrogels were incubated in 1mL of 1x PBS containing 0.02% NaN_3_ at 37°C for 14 days. At each time stage (1, 3, 5, 7 and 14 days), the incubation solution was collected and replaced with 1 mL of fresh PBS.

To determine the release kinetics of VEGF from alginate microspheres, 370 ± 77 mg of microspheres were incubated in 500 μL of 1x PBS containing 0.02% NaN_3_ at 37°C up to 14 days. At each time stage (1, 3, 7 and 14 days), samples were briefly centrifuged to separate the microspheres from the incubation solution, and 200 μL of the incubation solution was collected for VEGF quantification. At each time stage, the collected aliquot was replaced with 200 μL of fresh 1x PBS + 0.02% NaN_3_. For either bulk hydrogels and microspheres, sampled incubation solutions (1-5 μL) were diluted (1:20 − 1:100) and analyzed with VEGF ELISA kit (Ray Biotech, USA). Sample absorbance was read at 450 nm with a Cytation™ 3 Plate Reader (Biotek) and protein content was compared to a VEGF standard curve (0 − 6 ng/mL). The resulting data were analyzed as a cumulative release over the time stages. For microsphere, the measured VEGF concentration was normalized to the weight of the microsphere samples.

### 2.6 Matrigel® tube formation assay

Following manufacturer protocol, 50 µL of Corning Matrigel® Matrix stock solution was pipetted into each well of a 96 transwell plate and incubated at 37°C for 60 minutes to gel. A cell suspension of 20,000 iCell® human iPSC-derived endothelial cells (P4) in 50 µL of “test medium” was then added in each well, and 10 mg of microspheres were placed into the transwells. The test medium was composed of Lonza endothelial basal medium (EBM) containing the following components of the EGM™-2 BulletKit™: FBS, hydrocortisone, R3-IGF, ascorbic acid, hEGF, Gentamicin-Amphotericin and heparin. Test medium lacked VEGF and bFGF, the two major activators of angiogenesis.

The plates were incubated at 37°C for 24 hours in a 5% CO_2_ atmosphere. The tested conditions included Alg, S-Alg and H-Alg unloaded or loaded with VEGF (low= 0.5 µg/ml or high= 4 µg/ml of 1% w/v alginate solution). Samples with no microspheres in the transwell inserts cultured in test medium, complete Endothelial Growth Media-2 (EGM2, containing all the components of the BulletKit™, including VEGF and bFGF), or complete EGM2 media added with 100 µM suramin to inhibit angiogenesis were used as controls. Bright field images of the networks were captured at several time points (1h, 4h, 8h, 18h, 24h) with a Nikon Eclipse Ti-E inverted microscope at 4x and 10x magnification. Network masks were obtained by automated object analysis on CellProfiler [26] or by manual segmentation of networks in the case of low-contrast images. The extent of endothelial network formation was quantified by the analysis of skeletonized masks using the Fiji plugin “Analyze Skeleton” [27].

### 2.7 Aortic ring assay

All animal work was carried out in strict accordance with the recommendations in the Guide for the Care and Use of Laboratory Animals of the National Institutes of Health. The protocol was approved by the Brown University Institutional Animal Care and Use Committee (Protocol Number 1702000256). Thoracic aortas were collected from Sprague Dawley male and female rats (200 − 300g). Following excision, the aortas were washed in serum free EBM2 medium and the lumen was flushed to remove the blood using a 27-gauge needle. Each aorta was cut into rings (∼1 mm thick) using a scalpel blade and was subsequently transferred to a 10 cm^2^ dish containing 5 mL of serum-free EBM2 with 1% penicillin/streptomycin and incubated at 37°C overnight. In a 24-well culture plate, 180 µl of collagen (1 mg/mL) mixed with 18 mg of microspheres made of alginate or chemically modified alginate were added as a small drop in the center of each well. The aortic rings were positioned into the collagen/alginate microspheres drop and the plate was then incubated at 37°C for 60 minutes to allow collagen gelation. Following incubation, the plate was fed with 1 mL of test medium. Medium was changed at day 4, and the experiment was concluded at day 7. Alg, S-Alg and H-Alg unloaded or loaded with VEGF (low= 0.5 µg/µl or high= 4 µg/µl) were tested in the aortic ring assay. Control rings were prepared in collagen gels without microspheres and were cultured in test medium, EBM2 or complete EGM2 (controls). Texas Red® labeled Lycopersicon Esculentum (Tomato) lectin (0.02 mg/mL in PBS) was added to the rings at day 5 to selectively visualize endothelial cells.

Bright field, fluorescent and confocal images of the aortic rings were captured across several days (1d, 3d, 5d, 7d) with a Nikon Eclipse Ti-E inverted microscope and Olympus FV-1000-MPE Multiphoton Microscope. Assessment of cell migration distance and 3D reconstructions of the developing endothelial networks were performed using Fiji and IMARIS software.

### 2.8 Statistics

Statistical analysis was performed using one-way ANOVA with Tukey post hoc analysis. p values are indicated where significant (p<0.05).

## 3. Results

### 3.1 Synthesis of modified alginates by sulfation and EDC chemistry

Sulfated alginate was created by the substitution of sulfate groups at C-2 and C-3 of mannuronic and guluronic acid monomers using chlorosulfonic acid, while heparin was coupled to the carboxylic acid groups of the alginate backbone by use of EDC and EDA (Figure 1A). FT-IR analysis on unmodified and chemically modified alginate powders after extensive dialysis demonstrated the covalent bonding of sulfate and heparin moieties on the alginate backbone. The appearance of the characteristic sulfate peak at 1220 cm^−1^, attributed to S=O stretching [21, 28], was observed for both S-Alg and H-Alg (Figures 1B and C). In the S-Alg spectrum, the modifications of minor peaks observed around 800 cm^−1^ can be attributed to S–O–C stretching and suggest the sulfation of uronic acid monomers of alginate (Figure 1B). As discussed by Ronghua et al. [21], the new peak appearing after sulfation at 1690 cm^-1^ in S-Alg (stretching vibration of the carbonyl groups of –COOH), indicated by the arrow in Figure 1B, is associated to the formation of carboxyl groups (COOH) after dialysis (dialysis buffer pH=6.8). As shown in Figure 1C, the H-Alg spectrum showed the main absorption band of sulfate at 1220 cm^−1^, and a newly emerging peak at 980 cm^-1^, attributed to the asymmetric stretching of S-O-C [29], suggesting that heparin was successfully conjugated to the alginate backbone via carbodiimide chemistry. The amount of alginate-bound heparin detected with the toluidine blue colorimetric assay after chemical modification was 200 μg heparin/mg of alginate.

**Figure 1.**
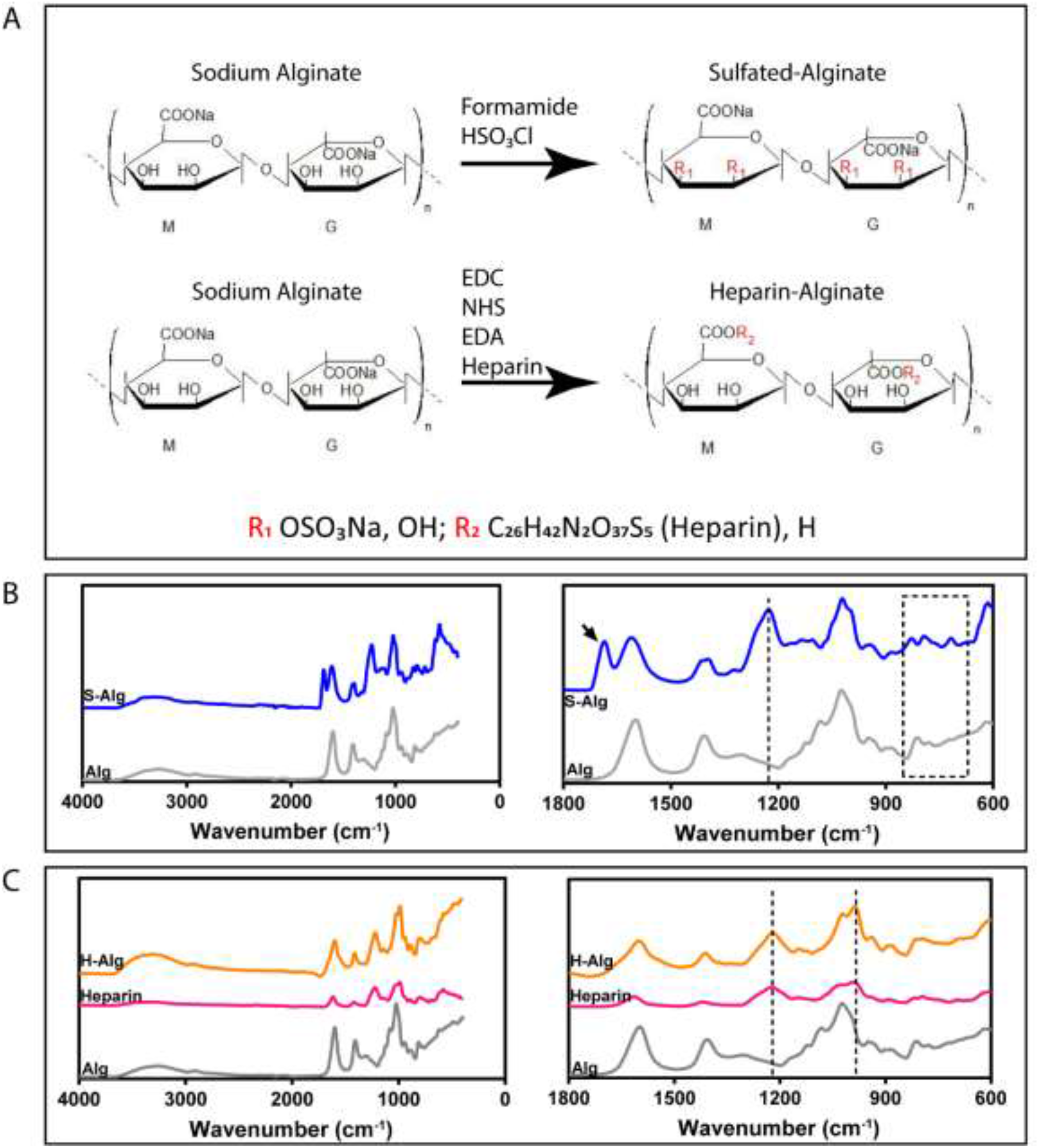
Bonding of sulfate groups and heparin moieties on alginate backbone. A) Scheme of the chemical modifications showing sites R1, R2 where modifications to sulfate (top) or heparin (bottom) occur to replace OH or H, respectively. FTIR analysis on S-Alg (B) and H-Alg (C) lyophilized powders show new major sulfate peaks at 1220 cm−1 (S=O symmetric stretching), modifications of the ring vibrations around 700 – 1000 cm-1, and post-dialysis carbonyl groups at 1690 cm-1 in S-Alg (arrow).

### 3.2 Alginate modifications influence the release profiles of VEGF

Alginate and modified alginate bulk hydrogels and microspheres were produced by internal and external gelation, respectively. S-Alg and H-Alg were obtained adding 5 mg of sulfated alginate and 10 mg of heparinized alginate to 1 g of unmodified alginate powders, respectively (0.5% and 1% modified Alg), before gelation as often done to preserve native gelation characteristics. Bulk alginate hydrogels (0.5 mL, 15 mm diameter) and microspheres produced with S-Alg and H-Alg showed sustained release kinetics of VEGF over a period of 14 days and a more gradual release between day 7 and 14 compared to the unmodified hydrogels (Figures 2A-B).

Microspheres exhibited a narrowly dispersed diameter, averaging 46 ± 15 μm, 37 ± 9 μm and 42 ± 12 μm for Alg, S-Alg and H-Alg, respectively (N.S). The size dispersion of the different formulations was comparable, despite a small sub-population of microspheres with diameters in the range of 70-80 μm was observed for Alg and H-Alg formulations (Figure 2C). The consistency in sizing within and across formulations permits the assumption that the release kinetics of immobilized proteins can be normalized for each sample based solely on mass. In all cases, protein was lost during extrusion and washing despite efforts to minimize the length of these processes and the encapsulation yield for Alg, S-Alg and H-Alg microspheres resulted in 70.0% ±18.3%, 66.7% ± 31.7% and 80.6% ± 12.6% encapsulation, respectively.

**Figure 2.**
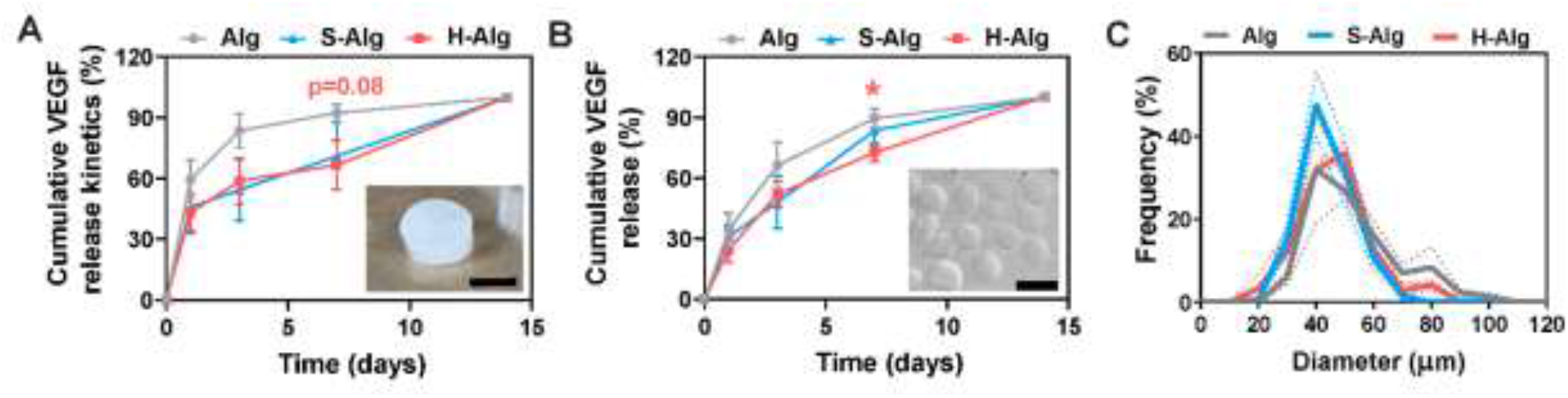
Production and release profiles of unmodified and chemically modified alginate hydrogels. A) Release profiles of VEGF from bulk alginate (Alg), sulfate-alginate (S-Alg) and heparin-alginate (H-Alg) bulk hydrogels (inset; scale bar: 1 cm), n=6. B) Release kinetics of VEGF loaded in Alg, S-Alg and H-Alg microspheres (inset; scale bar: 100 μm), n=5 (Alg and H-Alg) and n=6 (S-Alg). The p values reported in pink at day 7 show the level of significance between Alg and H-Alg groups (unpaired, two-tailed T-test, *p<0.05). C) Size dispersion, calculated by measuring the diameter of n=166 microspheres per formulation (Alg, S-Alg and H-Alg). Analyzed microspheres were pooled from 3 different production batches for each group. Solid line represents the mean, and dotted lines the SEM for each formulation.

In comparison to the VEGF release profiles reported for unmodified alginate microspheres in our previous work [30], where we have observed a burst 3-days release of biotinylated VEGF with western blot analysis, the sensitive ELISA assay used in this work allowed us to detect the release of small amounts of VEGF (5-20 ng/mL) in the incubation solutions at longer time points up to 14 days. The release kinetics of H-Alg bulk hydrogels and microspheres showed a more gradual release of VEGF over 14 days compared to the unmodified group, when normalized to total release at 14 days (Figures 2A-B). These results and data from other groups [23, 31] motivated an evaluation of the bioactivity and efficacy of the released VEGF with chemically modified alginate microspheres in translational *in vitro* and *ex vivo* assays of vascularization.

### 3.3 *H-Alg microspheres improve hiPSC-endothelial network formation* in vitro

A Matrigel® network formation assay was performed using human iPSC-ECs cultured in the presence of the microspheres, to assess the stability and proangiogenic activity of VEGF released from the microspheres. VEGF was loaded in alginate and chemically-modified alginate microspheres at low (0.5 µg/mL) and high (4 µg/mL) doses, to obtain a controlled release of the protein within the same order of magnitude of the concentration of VEGF added to the endothelial cell growth medium (2 ng/mL).

As shown in Figures 3 and 4, the chemically modified alginate microspheres loaded with VEGF produced either comparable or improved network formation with respect to the soluble delivery of multiple angiogenic factors (including VEGF, bFGF and IGF) from the complete EGM2. No microvessels were formed in the test medium (EGM2 without VEGF and bFGF) or in the samples treated with 100 µM suramin (negative control, data not shown). Increasing VEGF dose resulted in increasing trends of network structure development with unmodified alginate and sulfated-alginate microspheres. Interestingly, H-Alg microspheres showed pro-angiogenic signaling in the absence of or with low-doses of VEGF, and there was no additional benefit with high dose VEGF. Heparin-bound alginate microspheres releasing 0.5 µg/mL VEGF showed the maximum positive impact on hiPSC-ECs morphogenesis, measured as the area occupied by endothelial cell networks (Figure 3A).

**Figure 3.**
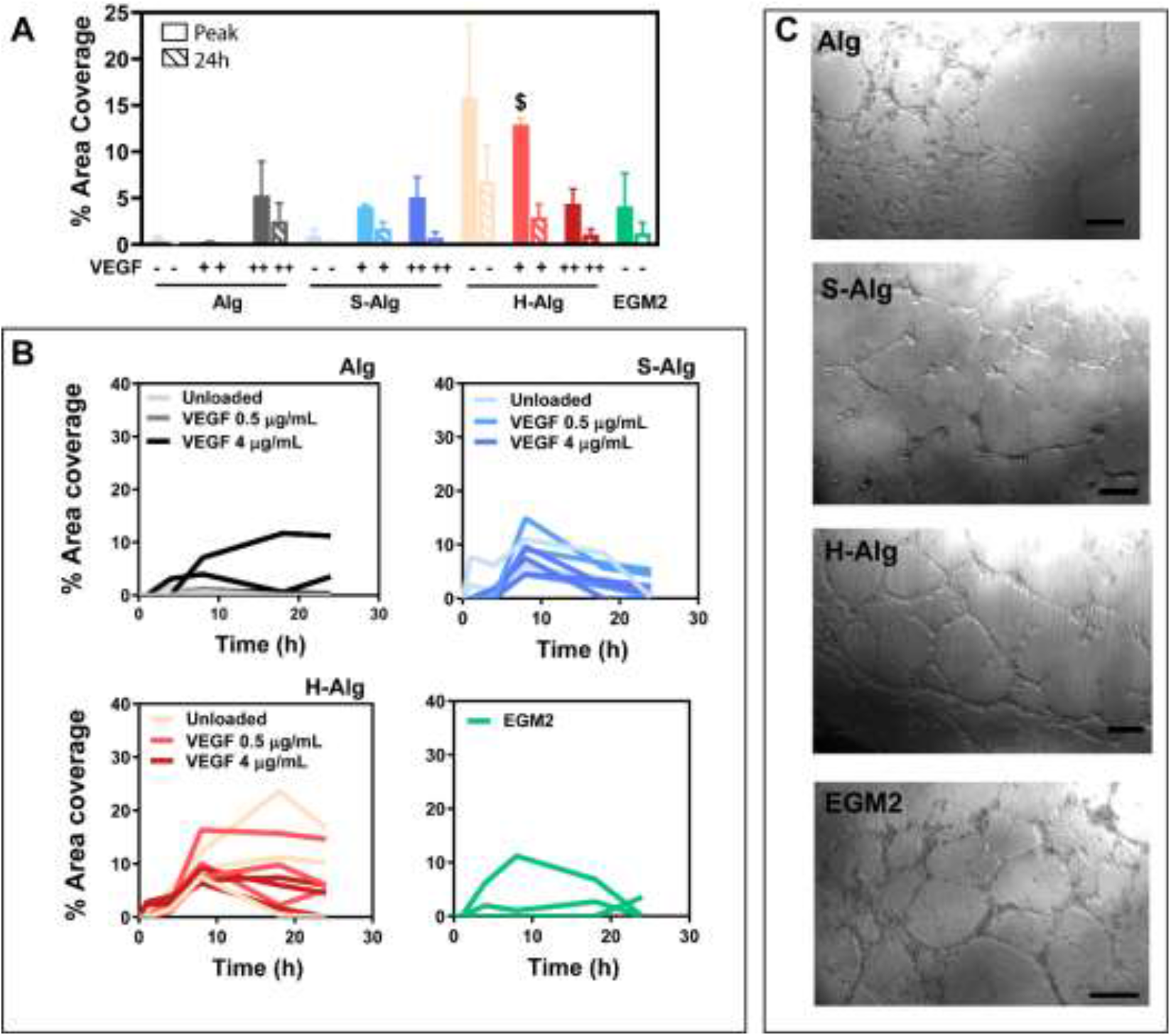
VEGF bioactivity and endothelial cell morphogenesis are enhanced with heparin-alginate microspheres in the network formation assay with defined collagen matrix. A) Measure of percent area coverage at the peak of network formation (reached between 8h and 18h for each sample) and at 24h hours. The graphs show the area of iPSC-endothelial cells networks treated with Alg, S-Alg and H-Alg microspheres loaded with 0.5 µg/mL (VEGF+), 4 µg/mL (VEGF++) or unloaded (VEGF-). The $ symbol indicates a significant enhancement in peak angiogenesis in the presence of heparin-modified alginate microspheres with respect of EGM2, Alg and S-Alg microspgheres loaded with 0.5 µg/mL VEGF (P<0.05). B) Quantification of network area over 24h shows the extent and kinetics of hiPSC-endothelial network formation in Matrigel® gels containing unloaded and loaded alginate microspheres. Each line represents a single sample. C) Images of networks developing in Matrigel containing unmodified alginate, sulfate-alginate, heparin-alginate microspheres loaded with 0.5 µg/mL of VEGF and EGM2 control. Images shown at 8 hours, scale bars = 200 µm.

**Figure 4.**
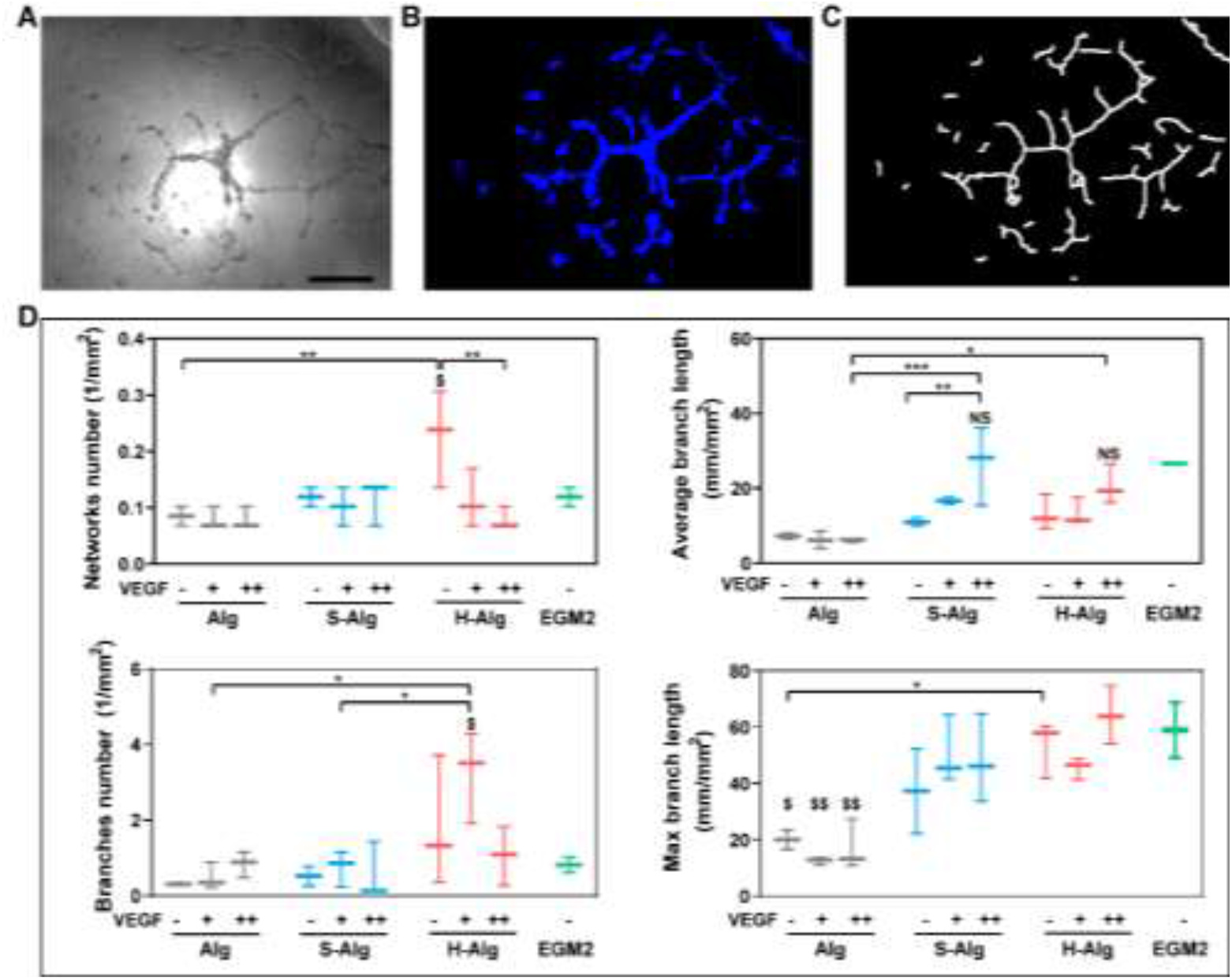
H-Alg microspheres enhance vessel-like structure formation compared to unmodified and S-Alginate microspheres as assessed by quantification of multiple angiogenic metrics in the hiPSC-derived EC network formation assay. Bright field images of networks are acquired at different times during the experiment, and n=3 images of different samples are analyzed for each group. The example in A) represents a H-Alg VEGF 4 μg/mL sample at 8h; scale bar = 500 µm. B) Network area mask obtained with a custom Cell Profiler code shows endothelial cell networks labeled with the blue color. C) Skeletonization of the networks with Fiji software finds the midline for quantitative analysis. D) Quantification of number of networks, average branch length, maximum branch length and number of branches. Tukey post-hoc test identifies significance (*P<0.05, **P<0.01 and ***P<0.001). NS indicates no statistical difference (P>0.05) compared to the EGM2 positive control, whereas $ denotes a statistical difference (P<0.05) vs. EGM2 control. $$ indicates a statistically significant relationship (P<0.05) with respect to both EGM2 control and the chemically modified alginate microspheres loaded with the corresponding concentration of VEGF.

The number of networks, average branch length, maximum branch length and number of branches were quantified to indicate network complexity and robustness of the angiogenic response in hiPSC-ECs at the peak of network formation (Figures 4A-4C). Similar to the network area coverage, the number of developing networks and network branches increases with H-Alg microspheres to a varying extent in unloaded microspheres or loaded with low (0.5 µg/mL) dose of VEGF, compared to Alg, S-Alg, and the EGM2 positive control (Figure 4D and Supplemental Figure 2). Furthermore, results indicate a significant increase in average and maximum branch length for samples treated with chemically modified alginate microspheres with respect to their unmodified counterparts. In the S-Alg and H-Alg samples, high VEGF dosage (4 μg/mL of 1% alginate solution) is determinant for achieving an average length of the developing networks comparable to that of the complete EGM2 control (Figure 4D). Based on these results, we have identified H-Alg microspheres as the most efficacious formulation for inducing vascular morphogenesis in hiPSC-ECs.

### 3.4 High dose VEGF released from heparinized alginate microspheres enhance angiogenesis kinetics in the aortic ring assay

The aortic ring assay is an *ex vivo* model that enables evaluation of the pro-angiogenic activity of released biologics in a complex, heterocellular environment [32] by following the formation and extent of network-like outgrowth from an aortic ring segment embedded in hydrogel. We have controlled the hydrogel chemical composition by using rat tail collagen type 1 and embedding microspheres, as opposed to a traditional Matrigel® hydrogel substrate. A significant increase in cell migration distance was observed at 1 and 3 days for samples containing H-Alg microspheres loaded with 4 µg/µL VEGF versus the EGM2 positive control (p<0.05), suggesting amplified cell migration for faster invasion of the collagen gel by native vascular wall cells (Figure 5A and 5B). Native cells remodeled the collagen matrix during cell migration and network formation, most notably in H-Alg and S-Alg samples, where migrating cells appear to pull on the collagen hydrogel to delaminate it from the aortic ring itself after 5 to 7 days (Figure 5A, red arrows). Detection of endothelial cells via fluorescently tagged lectin confirmed the presence of dense microvessel-like structures in the collagen with heparin-modified alginate microspheres scaffolds (Figure 5C).

**Figure 5.**
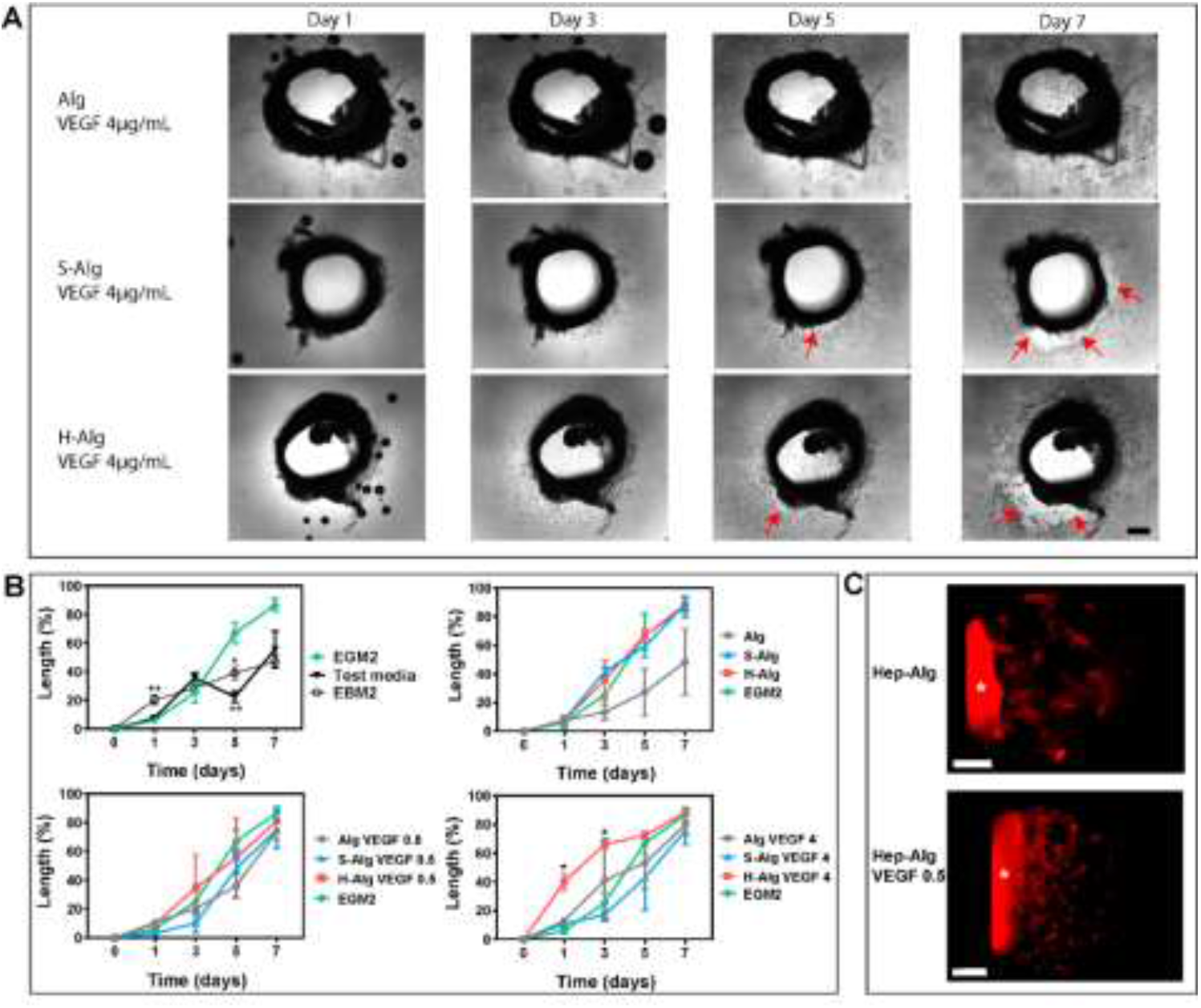
Native cell outgrowth kinetics from aortic rings and formation of 3D microvascular networks are improved with 4 µg/mL VEGF released from Heparin-Alginate microspheres. A) Native cells migrate from the aortic ring and invade the collagen scaffolds with embedded microspheres. The round black circles visible in the images at days 1 and 3 are air bubbles formed on the surface of cell culture medium. Red arrows point to sites where collagen delaminates from the aortic rings. Scale bar= 500 µm. B) Migration of the aortic ring cells into the collagen-alginate microspheres gel is assessed by measuring the outgrowth length in 2D (normalized by the total length available for outgrowth in 8 different locations around the ring). Panels of Figure 5B represent the investigated formulations: controls (top left), unloaded microspheres (top right) and microspheres loaded with 0.5 µg/mL VEGF (bottom left) and 4 µg/mL VEGF (bottom right). The same curve representing the EGM2 positive control is shown in each panel for comparison. * P<0.05, **P<0.01 compared to EGM2. C) Representative images of aortic rings labeled with Texas Red® labeled Lycopersicon Esculentum (Tomato) lectin at 7 days. Lectin binds to endothelial cell networks outgrowing from the rings and organizing into 3D structures. The white asterisk indicates the aortic ring. Scale bar= 500 µm.

## 4. Discussion

Revascularizing wound beds to regenerate functional tissue, such as skin ulcers in diabetic patients or the heart wall after myocardial infarction, requires customized approaches to locally instruct the microvasculature. Our approach uses alginate microspheres for hyperlocal delivery of growth factors that is amenable to integration with engineered tissues and for stabilizing proteins to extend the biological half-life by providing 1) heparin and sulfate groups for high-affinity binding (and therefore protection from proteolytic enzymes) and 2) physical entrapment into alginate hydrogels. Importantly, we demonstrate the efficacy of this biomaterial system using translationally relevant human iPSC-derived endothelial cells *in vitro*, an increasingly translational cell type for vascular tissue engineering, and a heterocellular primary rat culture *ex vivo* of aortic vascular cells. Sulfate groups and heparin are highly prevalent in the extracellular matrix, where active, secreted VEGF is involved in vasculogenesis (the appearance and coalescence of vessels) and angiogenesis during embryonic development [33] and in endothelial cell survival and wound healing in the adult [34, 35]. To mimic the strong interactions between heparin and VEGF (K_D_ = 40-157 nM) [36, 37], we have covalently bound alginate with sulfate groups or heparin moieties. Although we have used a lower density of sulfate and heparin groups for the production of modified alginate microspheres compared to other groups [23, 28, 38] with 0.05 sulfate per alginate monosaccharide and 20% heparin content in H-Alg, we found enhanced bioactivity of the immobilized VEGF *in vitro* compared to the unmodified alginate samples on hiPSC-ECs (Figures 3 and 4). As shown by others [22, 23, 28], sulfation and heparinization reactions impaired alginate ability to form microspheres via external gelation, making it necessary to mix unmodified alginate powders with 5 mg S-Alg/g of alginate or 10 mg H-Alg/g of alginate for optimal microspheres homogeneity and size distribution. To the best of our knowledge, in this work we demonstrate for the first time the proangiogenic effects of sulfated and heparinized micrometric-sized alginate microspheres (average size of 40 − 50 μm, Figure 2B), which is a convenient size for handling the microspheres and downstream therapeutic applications such as injecting them *in vivo* via small needles (25G − 30G) or for distributing them in soft hydrogels for applications in drug/growth factor delivery and tissue engineering [13]. We show that alginate heparinization offers advantages for revascularization therapy due to faster cell migration and network formation kinetics at critical early time points of 1 – 3 days (Figure 5) compared to unmodified alginate and the sulfated analog [39], likely due to the more specific binding interactions with the heparin-binding domain on VEGF for retention and protein preservation. We have also demonstrated for the first time a biomaterial-driven angiogenesis induced with unloaded heparinized alginate microspheres, where the modified biomaterial alone provided improved network metrics (Figure 4D), that we hypothesize is due to the localized heparin efficiently retaining autocrine and paracrine bioactive signaling molecules in the cellular microenvironment. Because too high and continuous levels of VEGF is known to result in aberrant angiogenesis and abnormal blood vessel growth leading to leaky and tortuous vessels [40], the loading dose of VEGF in microspheres was carefully calculated. We accounted for a 30% VEGF loss during extrusion and the half maximal release of 3-5 days (Figure 2), so that the daily amount of the protein released from the microspheres does not exceed the estimated physiological concentration range (32.3–70 ng/10^6^ cells per day [41]).

To further improve the relevance of our study to a heterogeneous cellular system as found in the vasculature of native tissue, we have evaluated our delivery system using complex native cell populations with a modified aortic ring assay in a chemically-defined collagen hydrogel. Host cell migration from rat aortic rings and 3-dimensional organization of endothelial network structures at 1 week provide evidence of the formation of stable networks (Figure 5). H-Alg microspheres loaded with 4 µg/mL VEGF resulted the most efficacious formulation for induction of angiogenic sprouting, showing improved efficacy versus the complete endothelial growth medium. With this work, we show that both endothelial network structure (Figure 3 and 4) and kinetics (Figure 5) are enhanced with hyperlocal controlled release of bioactive VEGF from small (40-50 µm) heparin-alginate microspheres, which are essential processes for effective vascularization therapy.

## 5. Conclusions

We have demonstrated that using a biomimetic binding interaction via heparin to prolong the bioactivity of a potent heparin-binding proangiogenic growth factor (vascular endothelial growth factor, VEGF) and control its release from alginate microspheres activates pro-angiogenic responses in complex, translationally relevant cellular systems with human iPSC-derived endothelial cells or native rat heterogeneous aortic vascular cells. The major benefit of increased kinetics of vascular network formations obtained via synergy between the heparin-modified biomaterial and growth factor-driven signaling suggests that this biomimetic system improves network morphology and may hasten wound healing and vascularization therapy, warranting advancement into tissue engineering applications.

## Acknowledgements

This research was supported by NIH R01 HL135091 (to KLKC) and by an Institutional Development Award (IDeA) from the National Institute of General Medical Sciences of the National Institutes of Health under grant number P20GM103652 (to FM). The authors are grateful to Fujifilm Cellular Dynamics, Inc. for providing the human induced pluripotent stem cell-derived endothelial cells used in this study.Dr. Anita Shukla for technical support in biomaterials characterization.

## Notes

### Competing Interest Statement

The authors have declared no competing interest.

