## Supplemental Figures for "Heparin-modified alginate microspheres enhance neovessel formation in hiPSC-derived endothelial cells and heterocellular *in vitro* models by controlled release of VEGF"

**Supplemental Material**

Supplemental Figure 1. Images of iPSC-derived endothelial cell networks at 24h in Matrigel® containing unmodified alginate, sulfate-alginate, heparin-alginate microspheres loaded with 0.5 µg/mL of VEGF and EGM2 control. Scale bars = 200 µm.

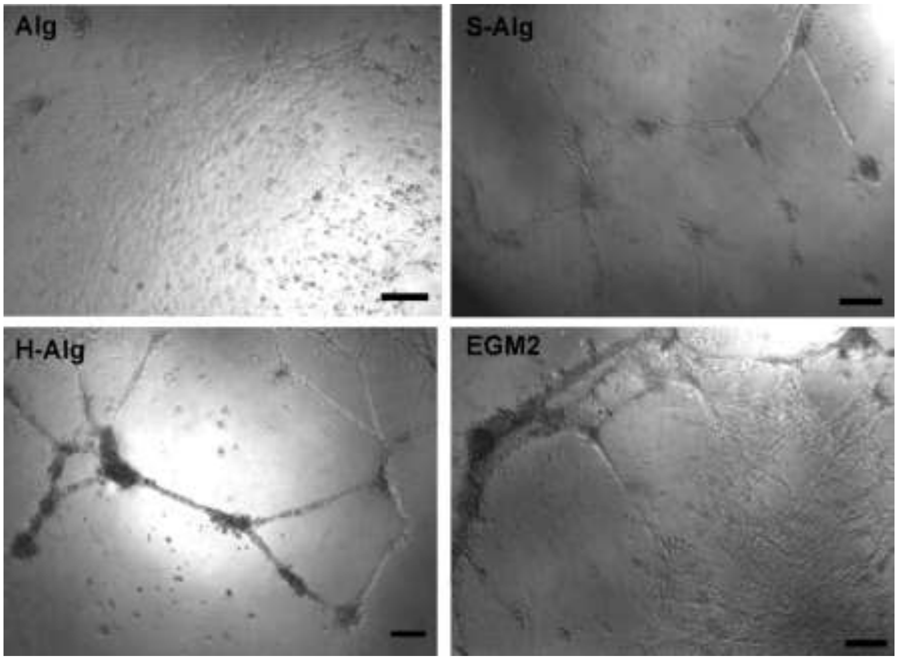

Supplemental Figure 2. Differences in morphogenesis of endothelial cells treated with heparinized versus sulfated alginate microspheres.

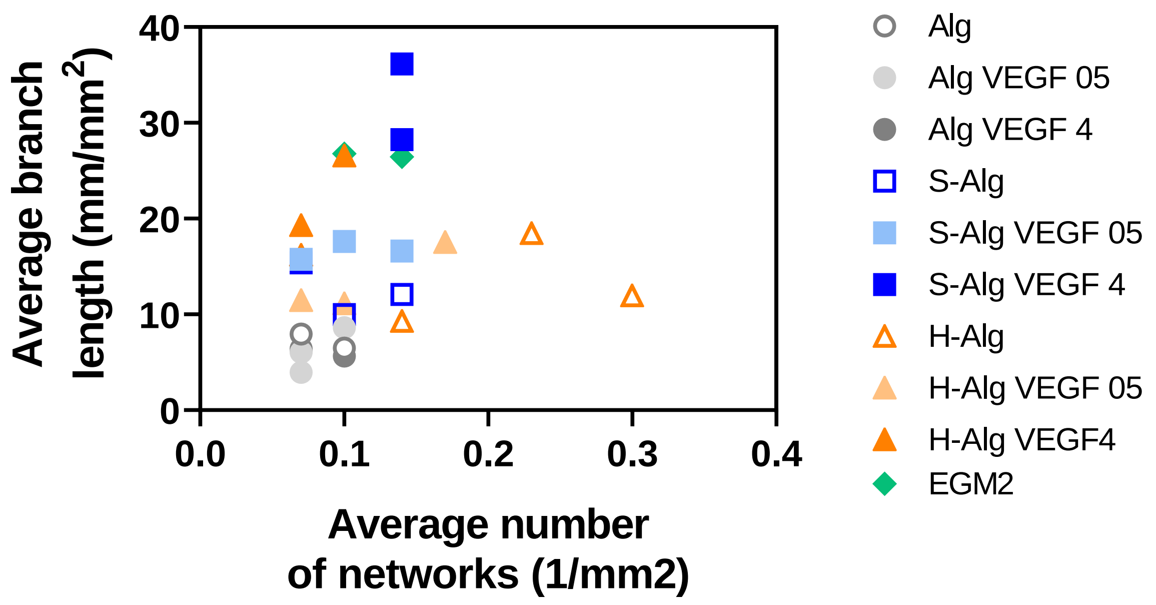
